# Dorsal striatum involvement in response conflict management – a lesion study in rats

**DOI:** 10.1101/2024.05.24.595791

**Authors:** Julien Poitreau, Boris Burle, Francesca Sargolini

**Affiliations:** Laboratoire de Neurosciences Cognitives, UMR 7291, CNRS, Aix-Marseille Université

**Keywords:** dorsal striatum, Simon effect, lesion

## Abstract

Action control allows to respond to relevant stimuli while ignoring the non-relevant stimuli in the surrounding environment. In humans this process is generally studied in conflict tasks, such as the Simon task, in which participants respond with a left or right button press to the non-spatial relevant feature (e.g. the color) of a lateralized stimulus, while ignoring the stimulus position. In this study we used a visual version of the Simon task that we have previsously developed in rats to investigate the involvement of the dorsal striatum, a brain area that is central in action control processes. We tested the effect of excitotoxic lesions of the dorsomedial (DMS) and dorsolateral (DLS) areas in learning to control response interference. We showed that both DMS and DLS lesions negatively impacted rat performances, and this effect strongly depends on task practice. These results suggest an involvement of both areas in learning to manage response conflict.

## 1 Introduction

Action control, among the cognitive control processes, is a mechanism allowing adapted behavior despite context instability or non-relevant interfering stimuli. It notably include appropriate action selection, incorrect responses correction, and conflict management between competing action plans (Ruitenberg et al., 2021). In humans, action control can be studied using the Simon task (Craft and Simon, 1967; Simon and Small, 1969), a conflict task in which participants are required to respond with a left or right button press to the non-spatial relevant feature (e.g. the color) of a lateralized stimulus, while ignoring the stimulus position. Although irrelevant to the task, the stimulus is presented lateralized, and can therefore define *compatible trials* when positioned on the same side as the required response (e.g. color-guided), or *incompatible trials* when positioned on the opposite side. Participants consistently display a decrease in performance in incompatible trials compared to compatible ones (i.e. the compatibility effect), resulting from a response conflict during action selection (Simon, Acosta and Mewaldt, 1975, Hasbroucq et al. 1999). The dual-route model (DRm - Kornblum, Hasbroucq, Osman, 1990) is classically used to explain it. This model assumes an automatic processing of the stimulus spatial feature, that leads to the activation of a stereotyped ipsilateral action plan, which interferes with the color-guided voluntary response in incompatible trials. Interference control mechanisms allows to resolve this conflict, but the process is time consuming and can be error prone, thus explaining the performance decrease.

The neural implementation of this psychological construct was found to rely on a well defined cortical network, including the frontal eye field (FEF), posterior parietal cortex (PPC) and dorsal premotor cortex (Bardi et al., 2012; 2015; Schiff et al., 2011), implicated in the automatic processing of the stimulus position, and a fronto-parieto-premotor network (Sebastian et al. 2013, Zhang et al. 2017), that is able to manage both the activation of the correct response and the inhibition of the automatic action plan in incompatible trials (Burle et al., 2016; van Campen et al., 2018). Moreover, several studies also support the involvement of the basal ganglia (BG - Fluchère et al. 2015; 2018; Sebastian et al., 2013; Schmidt et al., 2018; 2020; Stocco et al. 2017; van Wouwe et al. 2016; Wylie et al. 2010). In particular, the BG was proposed to be the site of the selection between the impulsive and controlled action plans (Wiecki & Frank, 2013), but this remains to be thoroughly investigated.

Although the BG function for spatial interference control remains relatively unexplored yet, other axes of research have long shown their involvement in processes conceptually close to what is described in the DRm. One of them, that is particularly influential, is the reinforcement learning theory for instrumental actions (i.e. the dual processes model in instrumental conditioning - DPm - Balleine, 2019), developed mostly in rodents. This theory also describes action control as a dual systems model, referred to as goal-directed and habitual systems, but centered on the dorsal striatum subregions (dorsomedial striatum - DMS and dorsolateral striatum - DLS, equivalent to the caudate nucleus and putamen in primates). Similarities between the DRm and the DPm are easy to apprehend, since both describe parallel systems competing for the response: a first system that is controlled, flexible, slow and resource consuming, and a second that is impulsive, stereotyped, fast and resource free (Dickinson & Pérez, 2018; Kornblum, Hasbroucq and Osman, 1990). So far, a single study has investigated the link between cognitive control and reinforcement learning processes (Otto et al. 2015). The results point to possible overlaps between the two processes, by showing a correlation between the ability to counteract impulsive behaviours in a Stroop task (i.e. another kind of conflict task - Stroop, 1935), and the tendency to use a goal-directed response strategy, in a two-step decision-making task in humans (Daw et al. 2011). To our knowledge, this question has never been addressed at the neural level. Given the central role of the dorsal striatum in the DPm, this structure appears to be a relevant starting point.

In this study we aimed at investigating 1) the basal ganglia involvement in action control under response conflict and 2) the overlap between cognitive control and reinforcement learning theories. To do so, we assessed the modifications in the acquisition and performance of interference control caused by specific excitotoxic bilateral lesions of the DMS and DLS, in rats performing a visual Simon task, that was validated in a previous study (Poitreau et al., submitted).

## 2 Methods and Materials

### 2.1 Animals

38 male Long Evans rats (Charles River Laboratories, Wilmington, USA) were used for this experiment. Animals were housed in pairs in a temperature- (21°C) and hygrometry-controled room, under a 12h dark-light cycle. All experimental procedures were done during the light phase. During the entire experiment, rats were food deprived and maintained at 85% of their free-feeding weight. They had unlimited access to water in their home cage. The rats included in this study were divided in two batches (batch 1: 8 SHAM, 3 DMS and 5 DLS; batch 2 : 8 SHAM, 8 DMS and 6 DLS), that were tested during two consecutive time periods separated by about 10 months due to the COVID-19 pandemic. All experimental procedures were approved by the local ethical committee and the French authority under the reference number APAFIS#9751-2017042712424200.

### 2.2 Simon task

The apparatus and the behavioral procedure are similar to those described in Poitreau et al. (sub-mitted).

#### Apparatus

Behavioural procedures took place in operant chambers (Med-Associates) of 29×24×29 cm (inner perimeter) with a grid as the floor, 2 side walls and ceiling in transparent acrylic glass, and front and back walls in stainless steel. The feeder (5×5×2 cm) was connected to a pellet dispenser, attached behind the back wall of the cage. The front wall was curved and contained 5 nose ports (NP), consisting in squares of 2.5×2.5 cm, 1 cm deep and positioned 2 cm above the floor. Only 3 NP were used, the central one and the 2 directly lateral on each side. The 2 others were blocked with a metal cap. All the NP were equipped with a light bulb inside, to provide visual stimuli, and infrared photocells allowed to detect the animals nose pokes. During the sessions, the chambers were completely in the dark, except for the light of the visual stimuli. A transparent acrylic wall with a 10×8 cm door was positioned 12 cm behind the front wall, and was used to control the position of the animals in front of the central NP. An infrared camera attached to the ceiling of the cage recorded animals behaviour. A loud-speaker attached to the ceiling of the cage provided auditory feedback (low-pitch or high pitch). Each chamber was in a sound-attenuating box, ventilated by a low-level noise fan. Cages were controlled by a computer and Med-Associates system, with programs wrote in Med-State Notation. Events during the task (i.e. stimuli, pokes, rewards) were recorded with Med-Associates system. Data were processed using MedPC-to-Excel, Python and R.

#### Response rule

Rats were trained to respond by a right or left nose poke in one of the two lateral NP according to the intensity (dim or bright) of a light stimulus, pseudo-randomly displayed in one of the lateral NP. The rule was counterbalanced across animals: half learned to respond with a left poke to a bright stimulus and a right poke to a dim stimulus, and half learnt the opposite rule.

Although non relevant for the task, stimulus position engendered two types of trial: *compatible trials*, when the stimulus position mapped the expected response side, indicated by the light intensity; and *incompatible trials*, when the two were opposite (for the precise temporal organisation of trial events and the experimental procedure timeline, see figure 1A and B).

**Figure 1.**
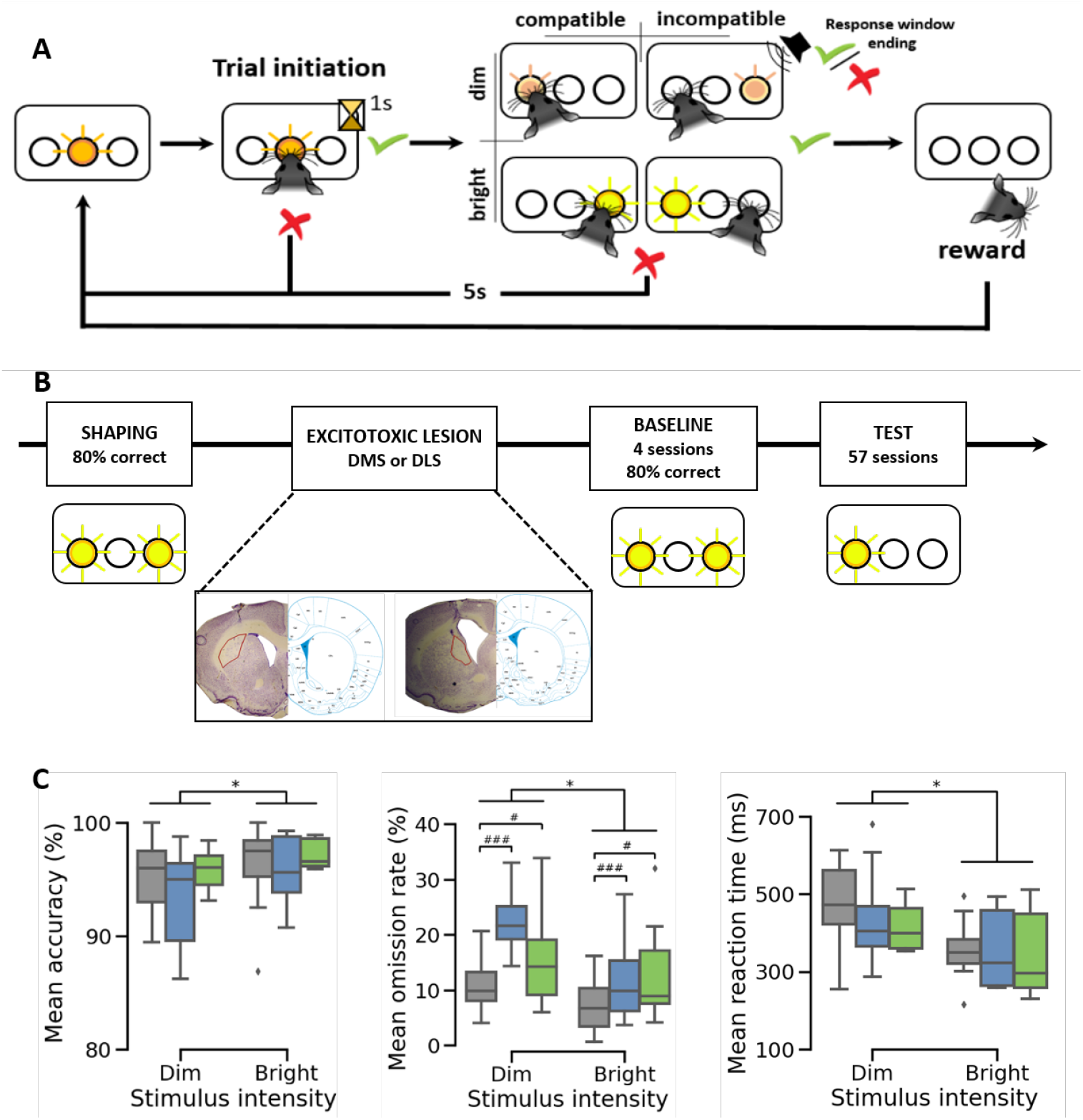
Behavioral protocol and baseline performance. **A**. Rat Simon task in a 3 nose ports set-up: The response rule for this example is “bright stimulus – right-side response” and “dim stimulus – left-side response”. A trial begins with a 1s nose poke in the central port that triggers the light stimulus (dim or bright). The response window is split in two periods: reaction time (RT – from stimulus onset to exit of the central port) and movement time (MT – from exit of the central port to entry into one of the lateral ports). The RT period is unlimited. The MT period is restricted to 800 ms. The end of the response window is indicated by an auditory feedback for response correctness. An incorrect response leads to a 5 s punishment delay with all the lights off. After a correct response, a sugar pellet is released following a poke into the feeder (located at the opposite side of the nose ports). **B**. Procedure timeline. Shaping: rule learning with bilateral light stimuli, before surgery. Baseline: Performance threshold assessment (*>*80% mean accuracy and *<*33% mean omission rate for both stimulus type during 4 sessions) after post-surgery re-shaping with bilateral stimuli. Test: 57 sessions with unilateral stimuli. **C**. Mean accuracy (left panel), mean omission rate (middle panel) and mean RT (right panel), in the 4 post-surgery baseline sessions, for the 3 groups (SHAM, DMS and DLS), and the 2 stimulus types (dim and bright). *, p*<*.05 significant effect of stimulus type (dim, bright); #, p*<*0.05 and # # #, p*<*0.001 significant effect of groups (SHAM, DMS and DLS).

**Figure 2.**
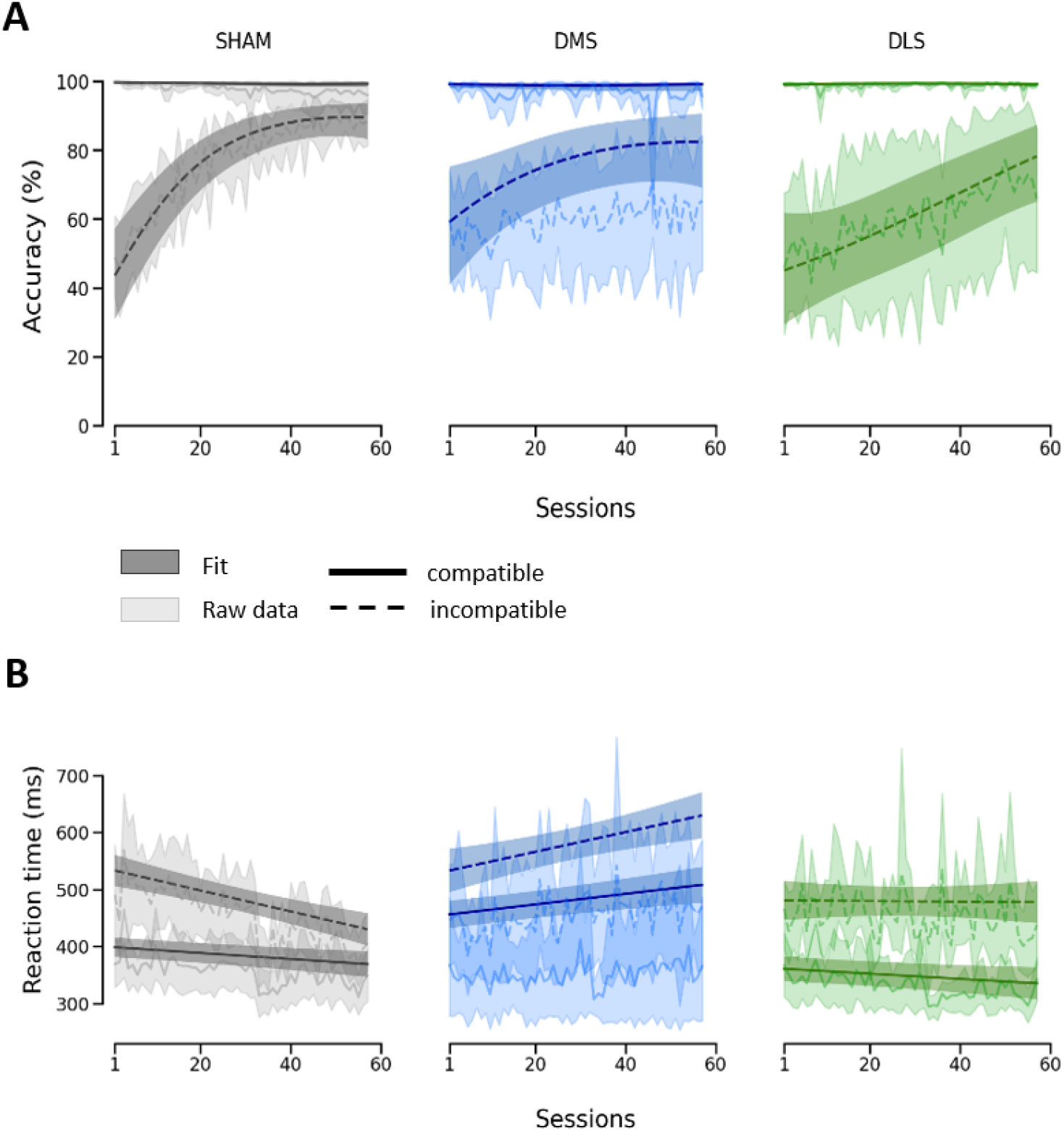
Mean performance across test sessions. Raw data and GLMM fits for accuracy (**A**) and reaction time (**B**) across sessions. The diagrams display mean and 95% confidence intervals for both raw data (light colors) and fit (dark colors).

#### Behavioural procedure

During all phases of the behavioral procedure, rats were trained in 30 minutes sessions, 2 times per day. The **pre-surgery shaping procedure** started with 4 sessions during which rats learnt the association between the feeder and the reward. In each session they had access to 45mg sugar pellets (sucrose dustless, TestDiet®, St Louis, Missouri) each time their nose was detected in the feeder, for a maximum of 150 pellets in a session. The next step consisted in teaching rats to initiate a trial with a nose poke in the central NP. The central port was enlightened to indicate the possibility to make a response. During the first four sessions, the pellet dispenser was controlled by a poke in the central NP (no time limitation). Following this, the pellet was triggered by a nose poke in the feeder after the central NP. Once the rat made a response, the light in the central NP was switched off until pellet retrieval. When rats achieved a performance criterion of 30 initiated trials with systematic reward retrieval per session, they underwent the next phase. Rats learnt to poke in a lateral NP after the initial central poke in response to a particular light stimulus intensity. After the central poke, a light stimulus (either bright or dim) was presented bilaterally in the lateral NPs, indicating that a lateral poke was awaited. During the first 2-4 sessions the same stimulus (alternatively bright and dim) was presented during the session. Then, the two stimuli were pseudo-randomly presented in the same session. The response time window was dissociated into reaction time (RT: from stimulus onset to central NP exit), and movement time (MT: from central NP exit to lateral NP entry). The temporal precision of both measures was 10 ms. No time restriction was imposed on RT, but MT could not exceed 10 seconds. Stimuli were displayed until the end of the MT window or until a response was given. A correct lateral poke was indicated by a high-pitch tone, and allowed to trigger the dispenser by a poke in the feeder. An incorrect lateral poke was indicated by a low-pitch tone, and was followed by a 5 sec punishment delay, during which all NPs were off. No response after the 10 seconds MT window was counted as an omission and treated as an error. To ensure accurate measures of RT, a premature response was counted for RT below 50 ms and treated as an error. A preparatory period, during which the rats had to hold their nose in the central NP, was also added by progressively increasing the delay before stimulus display from 10 msec to 1 sec. Once the animals achieved the learning criterion (80% of correct responses and less than 33% of omissions for both stimulus intensities), the MT window was progressively reduced from 10 sec to 800 ms, in order to speed up the responses, and hence get closer to the human definition of the reaction time (i.e. the time needed to respond as fast and as accurate as possible). When rats displayed correct performance (above learning criterion) during 4 consecutive sessions, they underwent the surgical procedure. For the **post-surgery procedure**, animals were re-trained with bilateral stimuli until they reached a correct performance (80% of correct responses and less than 33% of omissions for both stimulus intensities) during 4 consecutive sessions. These 4 last re-training sessions were used a posteriori, during the behavioral analyses, as baseline performance to assess group differences before the test phase. Then, the test phase started, with stimuli presented unilaterally, thus introducing stimulus position interference. All rats underwent 57 test sessions.

### 2.3 Surgery

Rats were deeply anesthetized with a mixture of oxygen and isofluran (4% for the induction, then 1%) associated with a subcutaneous injection of opioid analgesic (buprenorphine, Vetergesic, 0.03mg/Kg) and placed in a Kopf stereotaxic apparatus (Kopf instrument, Tujunga, CA, USA). Bilateral excitotoxic lesions of the dorsal striatum sub-regions (i.e. medial or lateral) were performed by injections of ibotenic acid (10mg/mL, Tocris, Bristol, UK) dissolved in a mixture of sterile NaCl (0.9%) solution with 10% NaOH (final solution pH: 7.8), via a stainless steel cannula (23 gauge) and tubing connected to a 10µL Hamilton syringe and an automatic pump (Harvard Apparatus, Holliston, USA). Injection rate was 0.08µL/min followed by a 5min diffusion time. The stereotaxic coordinates of the injection sites were determined based on the atlas of the rat brain (5th edition, Paxinos G. and Watson C., 2004, Elsevier). The volume of the injections was 0.2µL. 11 rats received dorsomedial striatum lesions: 3 injection points in each hemisphere at the following coordinates (relative to the bregma): 1) AP = 2.04mm, ML = ±1.7mm, DV = -5mm; 2) AP = 0.96mm, ML = ±2.2, DV = -5.2mm; 3) AP = -0.36mm, ML = ±3mm, DV = -4.6mm. 11 rats received dorsolateral striatum lesion at 3 injection points: 1) AP = 2.04mm, ML = ±3mm, DV = -4.8mm; 2) AP = 0.96mm, ML = ±4, DV = -5.2mm; 3) AP = -0.36mm, ML = ±4.2mm, DV = -5.2mm. 16 rats were used as sham controls and received injections of sterile NaCl solution (0.9%) at the same coordinates (8 DMS and 8 DLS). Following surgery, all animals received subcutaneous injections of a broad-spectrum antibiotic (oxytetrocycline 10%, Vetoquinol, 10mg/Kg) and a nonsteroidal anti-inflammatory drug (carprofen, Rimadyl, 5mg/Kg) and were allowed 14 days of recovery.

### 2.4 Histology and lesion quantification

At the end of the experiment, rats were killed using a 1mL intra-cardiac injection of a pentobarbital solution (Euthasol, 390mg/mL). Brains were then removed and frozen in dry ice. 40µm-thick coronal slices were mounted on glass slides and stained with thionin or labeled with the mitochondrial marker cychrome-C-oxydase. Lesion size and position (AP and ML coordinates) were determined from digital pictures, acquired with a Leica Microscope (Wetzlar, Germany), using an automatic program (with manual refinement) we developed, using ImageJ (Schneider et al., 2012). Based on the size and position of the lesion in each coronal section, we estimated the individual lesion centroids in each brain hemisphere. The lesion extent was expressed as the ratio between the volume of the lesion and the volume of the dorsal striatum (including DMS and DLS). Histology is depicted in the supplementary materials. No statistical quantification was performed. Following this, 7 rats (3 DMS and 4 DLS) with bad lesion positioning (overlapping on both the DMS and DLS), were excluded from the behavioural analyses. 3 SHAM were also excluded due to mechanical lesion on one or both striatum.

### 2.5 Data analyses

1 SHAM rat was excluded from the analyses after the test phase, because it displayed only 11% of mean correct responses over the 57 test sessions. From the whole experiment, 27 rats in total were finally included: 12 SHAM, 8 DMS and 7 DLS. We used the 4 last pre-test sessions with bilateral stimuli as baseline, to assess the response rule knowledge, and the differences between groups in processing the stimulus without the interference. Baseline groups differences were evaluated on the mean accuracy (i.e. % of correct responses), omission rate and reaction time (i.e. the time from the stimulus onset to the central NP exit). Mean accuracy and omission rate data were arcsine square root transformed to stabilise their variances. Mean reaction times were log-transformed to improve normality of the data. Data were analysed using analyses of variances (with stimulus intensity as within-subject factor and group as between-subject factor).

For the tests (i.e. unilateral stimuli), the accuracy and the correct RT were used as dependant variables, and were measured independently for each compatibility condition. Responses with RT superior or inferior to 2 times the median absolute deviation of the RT distribution, for each animals, were considered outliers and were excluded from the analyses (3.5% of the data – Leys et al. 2013). Response omissions (12.7% of the data), as well as anticipations, defined as RT *<* 170ms (3.6%), were also discarded.

Accuracy and correct RT of the 57 test sessions were analyzed using generalized linear mixed models (GLMM – Bates et al. 2015; Lo and Andrews, 2015). Models were constructed in an incremental manner using the lme4 package of the R language (v1.1-3.1 – Bates et al., 2014). Numerical variables were scaled when necessary. Following Bates et al. (2018), parsimonious random effect structures were determined using principal component analyses, in addition to several tests and metrics (AIC, BIC, likelihood ratio test, Bayes factor). Normality and homoscedasticity assumptions of the models residuals were assessed using the DHARMa package (v0.4.6 – Hartig & Hartig, 2017) when possible, or by visual inspection of the residuals plots. Overdispersion, for binomial models, was estimated using the sum of squared Pearson residuals (Bolker et al., 2009), and accounted for by adding an observation-level random effect (Harrison, 2015). Finally, hypothesis tests on fixed effect predictors were done by calculations of the estimated marginal means over models predictions, with significance testing using Wald chi-square tests of the car package (v3.1-2 – Fox et al. 2012). When necessary, we also used post-hoc pairwise comparison over the marginal means (emmeans package v1.8.3 – Lenth & Lenth, 2018; ggeffect package v1.1.4 – Lüdecke, 2018) with appropriate p-values correction, using the multivariate t method for multiple comparison p-values adjustment.

Our study aimed at investigating the acquisition and performance of interference control across the 57 test sessions as well as the impact of the dorsostriatal lesions. To this end, we analysed the evolution of mean accuracy and correct RT with practice to the task for the two compatibility conditions, according to the lesioned-groups, and also extracted from this the modulation of the compatibility effect (i.e. the difference in performances between the compatible and incompatible conditions). Furthermore, we used distributional analysis of the RT to describe performance modulation by practice and lesions, according to the response length, a factor that was long shown to be correlated to interference control efficiency (De Jong et al. 1994; Ridderinkhof, 2002) and, as such, to impact the compatibility effect. We tested adding the lesion size and the lesions antero-posterior coordinates in our different models but it did not bring conclusive results. Therefore, these variables were removed from the final analyses. The models employed are detailed below.

#### Mean performances progression across sessions

We first quantified performances progression trend for the three groups in each compatibility condition. To decrease the computational cost of the modeling, data were averaged per session and trial type for each rat. The sessions were modeled as a numerical vector of 57 ordered points.

##### Accuracy

We used a logit function as the model link function in order to respect the binomial distribution of the accuracy data. A quadratic term was added for the predictor sessions, to account for the the non-linear progression of the data, which significantly improved the model fit (linear model: AIC = 13502, BIC = 13707, log-likelihood (loglik) = -6716.9; quadratic model: AIC = 11407, BIC = 11757, loglik= -5645.4; p*<*0.001). As fixed-effect predictors we included: a fixed intercept, the linear and the quadratic terms for the sessions, the compatibility conditions, the experimental groups and the batch of training, as a possible confounding factor that we needed to control for. All the pairwise interactions between these predictors were also included. The sessions and compatibility predictors varying within the hierarchical grouping factor (i.e. the rats), we thus considered them to be included in the random effects component of the model. After refinement of the random effect structure, we kept in the model a random intercept, the main effect of the compatibility condition and the main effect of the sessions linear term, as well as the interaction between these two factors, and the interaction between the compatibility and the quadratic term of the sessions. All correlations between the random effects factors were estimated.

##### Reaction time

Following Lo and Andrews (2015), we assumed that the RT followed an inverse gaussian distribution and used an identity link function. Visual inspection of the data suggested a linear progression of the RTs across sessions. We thus modeled them as such. We included as fixed-effect predictors a fixed intercept, the sessions, the compatibility conditions, the experimental groups and the batch of training as well as all their interactions. As random-effects, we included the random intercept, the main effects of sessions and compatibility conditions, the interaction between the two factors and all their correlations.

#### Reaction time distribution analysis across test phases

Interference control processes required in the visual Simon task were shown to be modulated by the response time length in both humans and rats (Simon, Acosta, Mewaldt and Speidel, 1976; De Jong et al., 1994, Poitreau et al., submitted). Moreover, in humans, this dynamic was shown to be highly impacted by pathologies affecting basal ganglia, such as Parkinson disease (Wylie et al. 2010). Therefore, dorsostrialtal lesions could be expected to influence these modulations of the compatibility effect in rodents. To evaluate it, we first simplified the dynamic across sessions, by defining three equal periods of 19 test sessions, corresponding to the early, middle and late test phases. We then analysed correct RT according to the distribution of response latencies in the three test phases, using cumulative density functions of the RT distribution (CDF). CDF of raw correct RT for compatible and incompatible trials of each test phases were calculated with a Vincent averaging (Ratcliff, 1979): for each rat, compatibility condition and phase, RTs were sorted in ascending order and binned into 7 quantiles of equal size. The means for each quantile were computed and then averaged per quantile across rats to obtain a “mean” CDF representative of the individual ones.

We next applied a GLMM on the CDF data, including all groups, compatibility conditions and test phases in a unique model. Quantiles were defined as a numerical vector of 7 ordered points. In order to capture the complex S shape of the CDF curves, we considered modeling the quantiles predictor as a polynomial of degree 2 or 3. Comparing goodness-of-fit of the different models using likelihood ratio tests showed advantage for the cubic effect model (linear model: AIC = 11774, BIC = 12176, loglik = -5806,.8; quadratic model: AIC = 10970, BIC = 11549, loglik= -5370.1; cubic model: AIC = 10490, BIC = 11250, loglik= -5094.1; p0.001). Fixed effects predictors of the model included a fixed intercept, the 3 polynomial terms for the quantiles, the compatibility conditions, the test phases, the experimental groups and the batch of training as well as all the interactions. As random effects, we included the random intercept, the main effects of the quantiles linear term and of the compatibility condition, the interaction between the quantiles and the compatibility, between the quantiles and the late phase of training, and the third order interaction between these three factors. Random effects were estimated as independent, as correlations did not improve model fit.

To illustrate elements of interest (i.e. the compatibility effect across groups and the effect of practice to the task), from this complex GLMM of 18 modeled curves, we calculated the delta-plots of the model fits (i.e. the difference between the compatible and incompatible CDF curves), and also traced the evolution of each CDF quantiles across test phases. Of note is that these graphs are purely illustrative. They show summaries of our CDF fits and are used to emphasize the influence of our predictors on the CDF data. Significance of the different effects reported were obtained by estimating them directly on the CDF fits, not on their summaries.

## 3 Results

### 3.1 Performance during baseline sessions

We first report baseline data - the 4 last sessions with bilateral stimulation, after surgery. The color-to-response side was well mastered since accuracy and omission rate were above the learning criterion for the three groups (*>*80% of correct responses and *<*33% of omissions - figure 1C). We also used these sessions to assess potential differences in stimulus processing between groups before the introduction of the spatial interference.

Animals performed better for the bright than the dim stimulus, with higher accuracy (F(1,24) = 5.68, p *<* 0.05), lower omission rate (F(1,24) = 8.42, p *<* 0.01) and faster reaction time (F(1,24) = 33.38, p *<* 0.001). The groups differed on the omission rate (F(2,24) = 8.31, p *<* 0.01), which revealed that both lesion groups committed more omissions than the SHAM rats, independently of the stimulus intensity (SHAM - DMS: t = -3.97, p *<* 0.01; and SHAM - DLS: t = -2.54, p *<* 0.05; instensity × groups: F(2,24) = 1.22, p *>* 0.3). We shall come back on this difference later.

### 3.2 Results overview

The following analyses aimed at investigating 1) the acquisition of the spatial interference control and 2) the impact of the dorsostriatal lesions over this dynamic. In a nutshell, the following reported results show that SHAM improved their performance during sessions, with an increased accuracy especially in incompatible trials, the compatible one being at ceiling, and a shortening of RT particularly on incompatible trials, leading to a decrease of the compatibility effect, that nonetheless never disappears.

Compared to SHAM, dorsostriatal lesions impacted the performance deferentially depending on the lesion locations. Lesion of the DMS completely abolished any learning, since the accuracy did not improve during sessions in incompatible trials (accuracy being at ceiling on compatible ones), and RT remained stable across sessions. Rats with DLS lesions increase the accuracy in incompatible trial (again, the performance was at ceiling in compatible trials). In addition, RT decreased across sessions in compatible trials, but did not in incompatible ones.

In the following paragraphs we will describe in more details the above summarized results.

### 3.3 Performance progression across test sessions

This first analysis of the test data emphasised the dynamic of mean performances with practice to the task. Raw data and GLMM fits are illustrated in figure 3.

**Figure 3.**
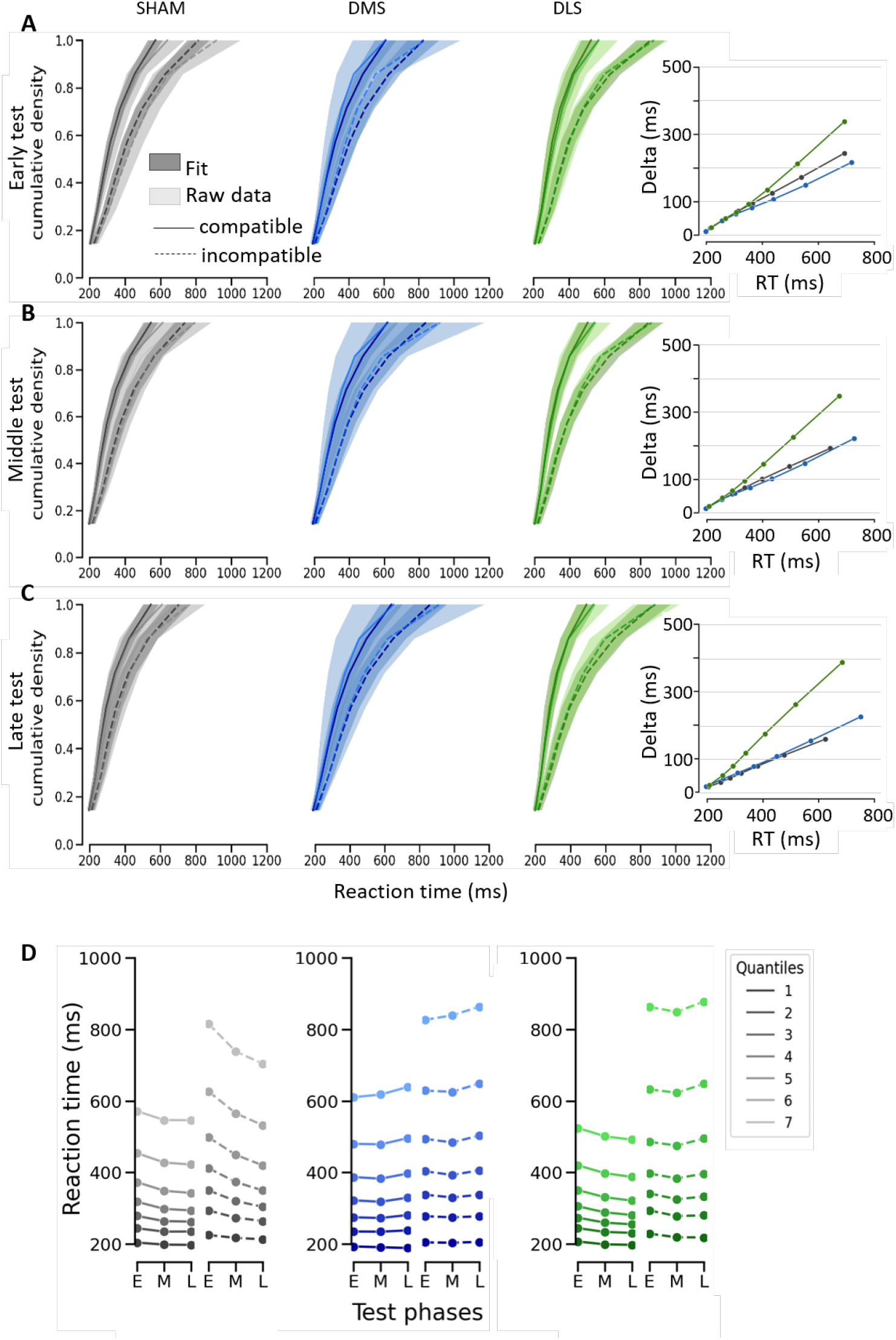
Cumulative density functions (CDF) of reaction times in correct compatible and incompatible trials, during the 3 test phases. **A**. Early training raw CDF (light colour) and GLMM fit (dark colour). **B**. Middle training raw CDF (light colour) and GLMM fit (dark colour). **C**. Late training raw CDF (light colour) and GLMM fit (dark colour). Graphs display mean and 95% CI of rats CDF curves and GLMM fit in each compatibility condition (compatible: full line; incompatible: dashed line). R^2^S= 0.978 was calculated for the entire model including the 3 groups, 2 compatibility conditions and 3 training phases

#### Accuracy

Correct response rate was higher on compatible than incompatible trials (χ^2^ = 277.25, df = 1, p *<* 0.001). Overall, accuracy increased with sessions (χ^2^ = 14.26, df = 2, p *<* 0.001), although this was modulated by compatibility (sessions × compatibility: χ^2^ = 38.2, df = 2, p *<* 0.001). This interaction between compatibility and session differed between groups (sessions × compatibility × groups: χ^2^ = 12.96, df = 4, p *<* 0.05).

More precisely, no change in accuracy was observed for compatible trials (at ceiling) in all three groups (SHAM: Marginal estimated trend - MET = -0.42, SE = 0.19, Z-ratio = -2.23, p = 0.14; DMS : MET = 0.007, SE = 0.21, Z-ratio = 0.032, p = 1; DLS: MET = 0.12, SE = 0.25, Z-ratio = 0.5, p = 0.99).

In contrast, for incompatible trials, SHAM and DLS rats significantly improved their accuracy across sessions (SHAM incompatible: MET = 1.42, SE = 0.23, Z-ratio = 6.21, p *<* 0.001; DLS incompatible:

MET = 0.88, SE = 0.28, Z-ratio = 3.13, p *<* 0.05), while DMS rats showed no improvement (p = 0.15). Accuracy improvement was not significantly different between SHAM and DLS (odds estimated ratio = 0.54, SE = 0.35, Z-ratio = 1.5, p = 0.65).

#### Reaction times

There was, overall, a clear compatibility effect, with shorter RT for compatible than incompatible trials (χ^2^ = 326.52, df = 1, p *<* 0.001). The three groups differed in their overall RT (χ^2^ = 52.72, df = 2, p *<* 0.001), but this difference depends on sessions (groups × sessions: χ^2^ = 18.96, df = 2, p *<* 0.001). The dynamic across sessions furthermore differed between groups and compatibility conditions (sessions × compatibility × groups: χ^2^ = 16.96, df = 2, p *<* 0.001).

Decomposing this evolution across sessions revealed that SHAM displayed a shortening of their responses in both compatibility conditions (compatible: MET = -17.6, SE = 5.3, Z-ratio = -3.32, p < 0.01 and incompatible: MET = -61.4, SE = 11.74, Z-ratio = -5.22, p *<* 0.001), but with a larger improvement in incompatible trials (compatible - incompatible estimated trends = 43.77, SE = 10.28, Z-ratio = 4.25, p *<* 0.001). In contrast to SHAM, DMS rats slowed down across sessions for both compatibility conditions (compatible: MET = 31, SE = 8.05, Z-ratio = 3.84, p *<* 0.001; incompatible: MET = 57.5, SE = 16.86, Z-ratio = 3.41, p *<* 0.01), at the same rate for the two (p = 0.08 for the difference in trend between the two conditions), although one has to remain cautious, since the fit was not very good Finally, DLS rats displayed no significant evolution of their RT across sessions, although there was a trend towards an improvement in compatible condition (compatible: MET = -15.2, SE = 6.51, Z-ratio = -2.331, p = 0.1; incompatible: MET = -1.8, SE = 15.07, Z-ratio = -0.119, p = 1).

### 3.4 RT distribution analysis

The previous analysis revealed the differential learning evolution across groups. To go one step further, we analysed RT distribution to study the dynamics of the underlying processes. However it is not realistic to analyse distributions of the 57 sessions. We hence divided the test sessions in three equal phases of 19 sessions, to assess averaged groups differences during the early, middle and late stages of practice to the task.

To investigate the influence of practice to the task, as well as the impact of the DMS and DLS lesions on this dynamic, we calculated CDF of the correct RT distributions with Vincent averaging (Ratcliff, 1979), which allows to draw curves representatives of the individual RT distributions. We then applied GLMM to describe and compare the shape of these curves across rats as a functions of groups and test phases. Data and fits are shown in figure 3, panel A, B and C. The dynamic of the GLMM fits as a function of practice is illustrated in the panel D. Delta plots of the GLMM fits - right-most panels of the figures 4A, B and C - illustrate the compatibility effect.

**Figure 4.**
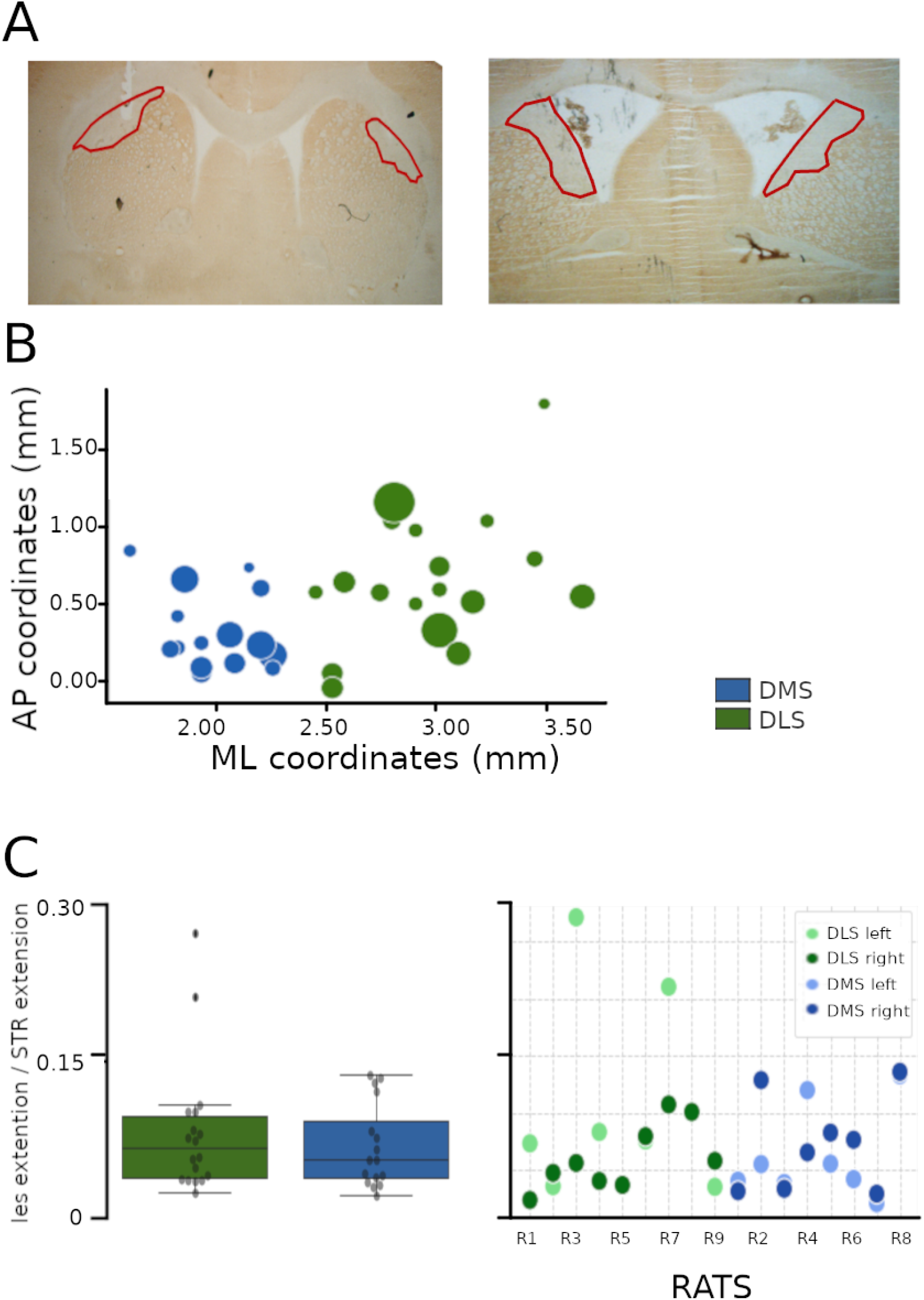
Histological verifications. **A**: Photos of coronal sections labeled with the mitochondrial marker cychrome-C-oxydase from two representative rats implanted in the DLS (left panel) and DMS (right panel). Lesions are circled in red. **B**: Scatter plot showing the antero-posterior (AP) and medio-lateral (ML) stereotaxic coordinates of the centroids of the lesions, in DLS (in green) and DMS (in blue) rats. Each circle represents the lesion centroid of one hemisphere. The size of the circle is proportional to the lesion extent. **C**: Left: box plots showing the normalized lesion extent (lesion size / dorsal striatum surface) for DLS and DMS rats. Right: Normalized lesion extent for each rats and each hemisphere.

The analyses displayed the overall main differences between groups (χ^2^ = 11.96, df = 2, p *<* 0.01) and between test phases (χ^2^ = 87.84, df = 2, p *<* 0.001). CDF shapes differed between compatibility conditions (quantiles × compatibility: χ^2^ = 345.27, df = 3, p *<* 0.001). This compatibility differences was also modulated by groups (quantiles × compatibility × groups χ^2^ = 29.93, df = 6, p *<* 0.001). Finally, test phases also influenced CDF shapes (quantiles × test phases: χ^2^ = 24.39, df = 6, p *<* 0.001).

To better clarify these effects, we analysed the learning effect on RT distribution shape for each group separately.

#### SHAM

SHAM animals speeded-up between the early and late phases, for both compatible and incompatible trials (See figure 3D, left panel), although it is more pronounced for incompatible trials. More specifically, speeding-up is present from quantile 2 to 7, p *<* 0.001 for all comparisons), while it is only present in a smaller portion of the RT distribution in compatible trials (from quantile 3 to 6, p *<* 0.01 for all comparisons). Regarding the compatibility effect, it was present on the entire RT distribution (i.e. from quantile 1 to 7 - p *<* 0.01 for all comparisons) and steadily increased (positive slope - p *<* 0.001). Moreover, the compatibility effect decreased in magnitude across test phases (early vs. middle phases from quantile 3 to 7: p *<* 0.05 for all; early vs. late phases from quantile 2 to 7: p *<* 0.05 for all), which is consistent with a faster decrease of the RT in compatible trials.

#### DMS

In the present analysis, no evolution in RT across test phases was evidenced for DMS rats (see figure 3D, middle panel - p *>* 0.1 for all quantiles), which is at odd with the data reported on section 3.3. The present analysis is performed on aggregated data, but on the whole RT distribution, and has a better fit. On the other hand, the previous analysis was aimed at capturing the whole learning curve, but only on means, and displayed a rather poor fit. Anyway, what is clear from these two analyses, is that the DMS rats did not improve their RT across session, hence presenting no learning. Despite this absence of learning, the compatibility effect (see delta-plots blue curves on figure 3, most-right panel) did not seem much affected. It was highly significant in all test phases, except for the shortest responses (quantile 1 early, middle and late: p *>* 0.05) and displayed a positive slope (apparently rather similar to the SHAM group). Moreover, it is not modulated across test phases, which is coherent with the constant response speed of this group, despite practice to the task.

#### DLS

As SHAM, DLS rats speeded-up in compatible trials, especially for the second half of the distribution (from quantile 3 to 6: p *<* 0.05 for all comparisons), a modulation that was only marginal on the mean analysis across sessions. No improvement was observed for incompatible trials with globally all p *>* 0.1, except between early and middle for quantiles 2 and 3 where p = 0.039 and 0.031. However, these difference are likely random variations, since they do not remain for the late phase. As for the other groups, the compatibility steadily increased (p *<* 0.001), and was highly significant for all quantiles (all p*<*0.001).

## 4 Discussion

The present study addressed the involvement of the two dorsostriatal sub-regions in the spatial interference control, as well as the possible links between the cognitive control and reinforcement learning theories formalized in the DRm and DPm. To this aim, we described the behavioural performances of rats with bilateral DMS or DLS lesions, while acquiring spatial interference control in a visual Simon task (Poitreau et al., submitted).

SHAM showed nearly perfect accuracy in compatible trials and a progressive improvement in the incompatible condition. They also improved RT in both compatibility conditions, with a greater RT decrease in incompatible trials. These effects lead to a long-lasting decrease of the compatibility effect in accuracy and RT. As for the lesion groups, DMS displayed greater deficits than the DLS, showing no improvement with practice in either accuracy or RT. Conversely, DLS lesion’s impact on performances is mostly visible in RT and appears to be compatibility dependent. Indeed, DLS accuracy is close to ceiling in compatible trials and improves in incompatible ones, similarly to SHAM. DLS also improved RT in compatible trials as the SHAM, but no improvement was observed in incompatible trials. This results in an increase of their compatibility effect on RT, unlike SHAM.

### 4.1 Practice effect for the control of spatial interference in rats and involvement of the dorsal striatum

Healthy rats improve their performance with sessions in both compatibility conditions. Baseline performance is correct, which suggests that this improvement is not related to the acquisition of the stimulus-response associations. Although, we evidenced baseline differences in performance between stimulus intensities and between groups in the omission rate, these parameters could not explain rat performances during the test. Indeed, stimuli were pseudo-randomly presented, to balance compatibility conditions and groups, and omissions trials were removed from the data. Overall, our experiment shows that the acquisition of the spatial interference control in rats leads to a decrease in the compatibility effect (both in accuracy and RT) that still persists after intensive learning (i.e. 57 sessions with an average of 86 trials each), similarly to humans (Proctor & Lu, 1999; Simon et al., 1973). Theses findings expands the results of our previous report showing similarities between the two species (Poitreau et al., submitted).

Regarding the dorsostriatal involvement, our results show no difference in performance between lesion and control groups in the early test phase. It thus suggests that action selection (correct or not) in both compatible conditions is independent of dorsal striatum when rats are not familiar with the spatial interference. In contrast, we show that practice improves both the efficiency and time course of the action selection, and this process is affected moinly by DMS but also DLS lesions. DMS group did not improve accuracy and RT during both compatible and incompatible trials, whereas DLS lesions exclusively prevented RT improvement in incompatible trials.

### 4.2 Relationship between the cognitive control and reinforcement learning processes

The DRm assumes that the Simon task brings into conflict two routes of information processing, automatic and controlled (Kornblum et al., 1990). Multiple studies support this view in both humans and rodents (e.g. Bardy et al., 2015; Burle et al. 2016; Ulrich et al. 2015; Poiteau et al. submitted) and suggest, in humans, the involvement of the basal ganglia in the process (Fluchère et al. 2015; 2018; Sebastian et al., 2013; Schmidt et al., 2018; 2020; Stocco et al. 2017; van Wouwe et al. 2016; Wylie et al. 2010). Our experiment allows to go further by demonstrating a differential involvement of the two dorsal sub-regions of the striatum in the improvement of the correct action selection in a situation of response conflict.

It seems important, however, to consider our results in a broader context than that of the DRm, by taking into account influential models of the striatal functions. The dorsostriatal dissociation is, indeed, extensively studied in the context of instrumental conditioning and it seems therefore reasonable to consider the DPm as one of the most influential model on the subject. According to this model (Balleine, 2019), and as shown by experimental data (Yin & Knowlton, 2004; 2005a), the posterior part of the DMS (pDMS) encodes response-outcome associations necessary for goal-directed responses, whereas the DLS encodes stimulus-response associations, at the core of habitual hehaviors. Goal-directed responses are, by definition, dependent on the response outcome (Dickinson & Pérez, 2018) and thus need to take into account explicit information such as the response rule. On the other hand, habits are by definition stimulus-based, automatic and outcome independent (Dickinson & Pérez, 2018).

The involvement of the dorsal striatum in the Simon task raises the question of the links between the DRm and the DPm. A simple hypothesis that can be advanced is that of a complete overlap between the two systems described. The controlled route may correspond to the goal-directed system aimed at developing an action plans based on the response outcome. The automatic route could be viewed as an overlearned (i.e. habitual) stimulus-response association between a spatially oriented cue and its ipsilateral response (see Umiltà & Zorzi, 1997 and Otto et al., 2015, for similar propositions). The strong deficits we observed in the DMS group are consistent with this view. Lesions of the DMS prevented responding in incompatible conditions, thus supporting an impairment of the controlled route. However, our results show that it is also the case for the DLS lesion, although to a less extent. From the DRm perspective, we can postulate that an impairment of the direct route should lead to a decrease of the compatibility effect, as we would have reduced the impact of the automatic response. DLS performances are therefore not in accordance with an overlap between the cognitive control and reinforcement learning processes described in the DRm and the DPm. Our results suggest that, in the context of the Simon task, goal-directed and habitual systems work both to select and execute the correct response in incompatible trials, and can therefore both be considered part of the controlled pathway.

### 4.3 Future directions

The links between cognitive control and reinforcement learning surely deserves to be more thoroughly explored, given the poor number of studies addressing the question - two to our knowledge: Otto et al. (2015) and ours - and the encouraging results obtained both at a cognitive (Otto et al. 2015) and neural level (the present study). Nonetheless, regarding the implementation of the spatial interference control at the dorsostriatal level, the functional dissociation between the DMS and the DLS seems not to be the ideal framework.

Our data could possibly be more easily explained according to the Wiecki and Frank (2013) model of a cortico-basal ganglia network for the resolution of conflict tasks. This model does not emphasise the DMS and DLS functional dissociation but rather the competition between the basal ganglia (BG) output patways (i.e. the direct and indirect pathways) as being critical for the correct action selection in both compatible and incompatible trials (see also Bariselli, Fobbs, Creed, & Kravitz, 2018). This model proposes that the action plans created during the Simon task are evaluated trial by trial inside the BG to allow their execution and the selection process in case of conflict. Cortical structures provide evidence for one or the other action plan by intervening on either the direct (GO) or indirect (NOGO) BG pathways encoding the two responses.

In this model, the goal-directed and habitual systems can be integrated as part of different cortico-striatal lopps (associative vs sensorimotor), that hence bring different information to favour one or the other action plan. Lesion of the DMS or DLS may thus remove the corresponding cortical information from the selection mechanism. The current literature shows that the integration of associative as well as sensorimotor information at the striatal level is correlated with performance progression in discrimination tasks (Bissonette & Roesch, 2015; Xiong et al. 2015). Accordingly, we show that both types of information are used to improve the selection of the correct action to perform in the Simon task. For both types of information, our results suggest that disruption of this integration tends to impair the possibility of selection improvement with experience, particularly in a situation of response conflict. We further showed that in such situation the system is more dependent on the associative process rather than the sensorimotor one. This result is consistent with Bissonette & Roesch (2015) study, which showed that the DMS is sensible to the response rule and to the conflict between potential responses, an information that seems particularly relevant to discriminate between several action plans. In future experiments it is crucial to explore the involvement of the direct and indirect BG pathways, in order to understand the action selection mechanism required by the spatial interference.

Another aspect that will be important to take into account is the heterogeneity of the cortico-striatal afferent (Hintiryan et al. 2016) and the functional dissociation of the striatal regions that result from it, notably inside the DMS. The DPm goal-directed system is based on the posterior part of the DMS since the anterior part is not necessary for goal-directed behavior (Yin et al. 2005b). Our work did not take into account this dissociation since the lesions spread along the entire antero-posterior axis (see supplementary material). Future works should aim at refining this aspect, in order to precisely determine cortico-striatal connections involved in the spatial interference control.

Finally, our results could be related to the different plasticity mechanisms occurring in the two dorsostriatal regions during learning (Hawes et al., 2015; Perez et al., 2022; Yin et al. 2009). Exploring them with respect of the compatibility condition as well as the striatal input and output could explain at the cellular level the behavioural results evidenced in the present study.

## Acknowledgments

We thank Kevin Poireau for his help on the histological analysis, Didier Louber and Dany Paleressampoule for technical assistance, the Ministry of the Research for fundings. J. Poitreau was funded by a Phd grant from the Ministry of Research, and a grant ‘en of PhD’ from the EUR Neuroschool.

## References

Balleine, B. W. (2019). The meaning of behavior: Discriminating reflex and volition in the brain. Neuron, 104(1), 47–62.

Bardi, L., Kanai, R., Mapelli, D., & Walsh, V. (2012). TMS of the FEF interferes with spatial conflict. Journal of Cognitive Neuroscience, 24(6), 1305–1313.

Bardi, L., Schiff, S., Basso, D., & Mapelli, D. (2015). A transcranial magnetic stimulation study on response activation and selection in spatial conflict. European Journal of Neuroscience, 41(4), 487–491.

Bariselli, S., Fobbs, W., Creed, M., & Kravitz, A. (2019). A competitive model for striatal action selection. Brain research, 1713, 70–79.

Bates, D., Mächler, M., Bolker, B., & Walker, S. (2014). Fitting linear mixed-effects models using lme4. arXiv preprint arXiv:1406.5823.

Bissonette, G. B., & Roesch, M. R. (2015). Rule encoding in dorsal striatum impacts action selection. European Journal of Neuroscience, 42(8), 2555–2567.

Bolker, B. M., Brooks, M. E., Clark, C. J., Geange, S. W., Poulsen, J. R., Stevens, M. H. H., & White, J.-S. S. (2009). Generalized linear mixed models: A practical guide for ecology and evolution. Trends in ecology & evolution, 24(3), 127–135.

Burle, B., van den Wildenberg, W. P. M., Spieser, L., & Ridderinkhof, K. R. (2016). Preventing (impulsive) errors: Electrophysiological evidence for online inhibitory control over incorrect responses: Inhibitory control of incorrect responses. Psychophysiology, 53(7), 1008–1019.

Craft, J. L., & Simon, J. R. (1970). Processing symbolic information from a visual display: Interference from an irrelevant directional cue. Journal of Experimental Psychology, 83(3, Pt.1), 415–420.

Daw, N. D., Gershman, S. J., Seymour, B., Dayan, P., & Dolan, R. J. (2011). Model-based influences on humans’ choices and striatal prediction errors. Neuron, 69(6), 1204–1215.

De Jong, R., Liang, C.-C., & Lauber, E. (1994). Conditional and unconditional automaticity: A dual-process model of effects of spatial stimulus-response correspondence. Journal of Experimental Psychology: Human Perception and Performance, 20(4), 731–750.

Dickinson, A., & Pérez, O. D. (2018). Actions and Habits. In Goal-Directed Decision Making (p. 1–25). Elsevier.

Fluchère, F., Burle, B., Vidal, F., van den Wildenberg, W., Witjas, T., Eusebio, A., Azulay, J.-P., & Hasbroucq, T. (2018). Subthalamic nucleus stimulation, dopaminergic treatment and impulsivity in Parkinson’s disease. Neuropsychologia, 117, 167–177.

Fluchère, F., Deveaux, M., Burle, B., Vidal, F., van den Wildenberg, W. P. M., Witjas, T., Eusebio, A., Azulay, J.-P., & Hasbroucq, T. (2015). Dopa therapy and action impulsivity: Subthreshold error activation and suppression in Parkinson’s disease. Psychopharmacology, 232(10), 1735–1746.

Harrison, X. A. (2015). A comparison of observation-level random effect and Beta-Binomial models for modelling overdispersion in Binomial data in ecology & evolution. PeerJ, 3, e1114.

Hart, G., Bradfield, L. A., & Balleine, B. W. (2018). Prefrontal Corticostriatal Disconnection Blocks the Acquisition of Goal-Directed Action. The Journal of Neuroscience, 38(5), 1311–1322.

Hartig, F., & Hartig, M. F. (2017). Package ‘DHARMa’. R package.

Hasbroucq, T., Possamäi, C.-A., Bonnet, M., Vidal, F. (1999). Effect of the irrelevant location of the response signal on choice reaction time: An electromyographic study in humans. Psychophysiology, 36(4), 522–526.

Hawes, S. L., Evans, R. C., Unruh, B. A., Benkert, E. E., Gillani, F., Dumas, T. C., & Blackwell, K. T. (2015). Multimodal plasticity in dorsal striatum while learning a lateralized navigation task. Journal of Neuroscience, 35(29), 10535–10549.

Hintiryan, H., Foster, N. N., Bowman, I., Bay, M., Song, M. Y., Gou, L., Yamashita, S., Bienkowski, M. S., Zingg, B., Zhu, M., Yang, X. W., Shih, J. C., Toga, A. W., & Dong, H.-W. (2016). The mouse cortico-striatal projectome. Nature Neuroscience, 19(8), Article 8.

Kornblum, S., Hasbroucq, T., & Osman, A. (1990). Dimensional overlap: Cognitive basis for stimulus-response compatibility–A model and taxonomy. Psychological Review, 97(2), 253–270.

Leys, C., Ley, C., Klein, O., Bernard, P., & Licata, L. (2013). Detecting outliers: Do not use standard deviation around the mean, use absolute deviation around the median. Journal of Experimental Social Psychology, 49(4), 764–766.

Lo, S., & Andrews, S. (2015). To transform or not to transform: Using generalized linear mixed models to analyse reaction time data. Frontiers in psychology, 6, 1171.

Lüdecke, D. (2018). ggeffects: Tidy data frames of marginal effects from regression models. Journal of Open Source Software, 3(26), 772.

Otto, A. R., Skatova, A., Madlon-Kay, S., & Daw, N. D. (2015). Cognitive control predicts use of model-based reinforcement learning. Journal of cognitive neuroscience, 27(2), 319–333.

Perez, S., Cui, Y., Vignoud, G., Perrin, E., Mendes, A., Zheng, Z., Touboul, J., & Venance, L. (2022). Striatum expresses region-specific plasticity consistent with distinct memory abilities. Cell Reports, 38(11).

Proctor, R. W., & Lu, C.-H. (1999). Processing irrelevant location information: Practice and transfer effects in choice-reaction tasks. Memory & Cognition, 27(1), 63–77.

Ratcliff, R. (1979). Group reaction time distributions and an analysis of distribution statistics. Psychological Bulletin, 86(3), 446–461.

Ridderinkhof, K. R. (2002). Activation and suppression in conflict tasks: Empirical clarification through distributional analyses. In Common Mechanisms in Perception and Action. Attention & Performance (Vol. 19, p. 494-519). W. Prinz B. Hommel.

Schneider, C. A., Rasband, W. S. & Eliceiri, K. W. (2012). NIH Image to ImageJ: 25 years of image analysis. Nature Methods, 9(7), 671–675.

